# Corticofugal oscillatory modulation of the cochlear receptor during auditory and visual attention in tinnitus

**DOI:** 10.1101/2023.09.20.558639

**Authors:** Rodrigo Donoso-San Martín, Alexis Leiva, Constantino D. Dragicevic, Vicente Medel, Paul H. Delano

## Abstract

The mechanisms underlying tinnitus perception are still under research. One of the proposed hypotheses involves an alteration in top-down processing of auditory activity. Low-frequency oscillations in the delta and theta bands have been recently described in brain and cochlear infrasonic signals during selective attention paradigms in normal hearing controls. Here, we propose that the top-down oscillatory activity observed in brain and cochlear signals during auditory and visual selective attention in normal subjects, is altered in tinnitus patients, reflecting an abnormal functioning of the corticofugal pathways that connect brain circuits with the cochlear receptor. **Methods:** To test this hypothesis, we used a behavioral task that alternates between auditory and visual top-down attention while we simultaneously measured electroencephalogram (EEG) and distortion-product otoacoustic emissions (DPOAE) signals in 14 tinnitus and 14 control subjects. **Results:** We found oscillatory activity in the delta and theta bands in cortical and cochlear channels in control and tinnitus patients. There were significant decreases in the DPOAE oscillatory amplitude during the visual attention period as compared to the auditory attention period in tinnitus and control groups, while frontal EEG oscillatory activity (Fz) was increased during the visual attention task only in the tinnitus group. The difference between auditory and visual DPOAE oscillatory power in the delta band was stronger in tinnitus individuals as compared to controls. **Conclusions:** These results confirm the presence of top-down infrasonic low-frequency cochlear oscillatory activity in the delta and theta bands in tinnitus patients. Our results show an increase in the EEG theta oscillatory power in frontal regions of tinnitus sufferers concomitant with a stronger corticofugal suppression of cochlear oscillations in the delta band during visual attention in tinnitus. These findings suggest an increased compensatory corticofugal inhibitory mechanism to the cochlear receptor in tinnitus.

## Introduction

Tinnitus is the perception of a phantom sound in the absence of external acoustic stimulation (Elgoyhen et al., 2015). It is a prevalent condition ranging from 11% to 30% of the general population, while for a subset of individuals, it becomes a chronic and distressful symptom (McCormack et al., 2016), affecting auditory, cognitive, and emotional brain networks (Husain 2016; De Ridder et al., 2021). Chronic tinnitus sufferers have a higher prevalence of neuropsychiatric conditions, mainly anxiety and depressive symptoms (Geocze et al., 2013; Ziai et al., 2017). In addition, cognitive abilities, such as selective attention, can be affected by tinnitus (Roberts et al., 2013), as the perception of the phantom stimulus could be perceived as an annoying distractor that constantly disrupts attention, altering performance in auditory and visual attention tasks (Araneda et al., 2015).

Although several hypotheses have been raised to understand the neurobiology of tinnitus, to date, the neural mechanisms of tinnitus are still elusive (Knipper et al., 2020). One of the proposed mechanisms involves alterations in top-down processing at different levels of the auditory pathway (Knipper et al., 2021). In this line, the auditory efferent system connects central auditory structures with cochlear hair cells and auditory-nerve neurons by multiple neuroanatomical feedback circuits (Elgueda and Delano, 2020) that might be altered in tinnitus sufferers (Knipper et al., 2020). The auditory efferent system is organized into (i) brainstem circuits (medial and lateral olivocochlear neurons) (Warr and Guinan, 1979), and (ii) corticofugal descending pathways connecting the auditory cortex with subcortical nuclei (Terreros and Delano, 2015; Lauer et al., 2022). Importantly, one of the functions that has been attributed to these corticofugal pathways is to suppress auditory responses during selective attention tasks (Oatman 1971; Delano et al., 2007; Srinavasan et al.,2012; Wittekindt et al., 2014; Terreros et al., 2016).

A possible alteration in the auditory efferent functioning in tinnitus patients has been studied by several researchers with contradictory results (Riga et al., 2015). However, it is important to highlight that previous studies searching for a role of the auditory efferent system in tinnitus have only studied brainstem circuits, specifically the activation of the medial olivocochlear reflex with contralateral acoustic stimulation (Riga et al., 2015). Thus, it is still unknown whether there is a functional alteration of the auditory efferent corticofugal projections during attention paradigms in tinnitus sufferers.

Recently, in non-tinnitus subjects, we described a top-down mechanism involving the corticofugal oscillatory modulation of cochlear responses at theta and delta bands (<10 Hz), during visual and auditory attention, using simultaneous electroencephalogram (EEG) and distortion-product otoacoustic emissions (DPOAE) recordings (Dragicevic et al., 2019). Furthermore, similar oscillatory corticofugal modulations have been observed in measurements of spontaneous external ear canal pressure (Kohler et al., 2021; Köhler and Weisz 2023) and auditory-nerve responses recorded with cochlear implants (Gemmacher et al., 2022), during visual and auditory selective attention tasks. Taken together, these independent works provide evidence of a top-down oscillatory mechanism in delta and theta frequency bands during selective visual and auditory attention that modulates cochlear and auditory-nerve responses.

Here, we propose that the corticofugal oscillatory activity observed during auditory and visual selective attention in normal subjects (Dragicevic et al., 2019; Kohler et al., 2021; Gemmacher et al., 2022) is altered in tinnitus patients, reflecting an abnormal functioning of the corticofugal pathways that connect the auditory cortex with the cochlear receptor. To test this hypothesis, we used the same attentional paradigm as in Dragicevic et al., (2019), simultaneously measuring EEG and DPOAE signals in tinnitus and control subjects.

## Methods

### Subjects

Subjects were recruited from the Otolaryngology Department of the Clinical Hospital of the Universidad de Chile, which were invited to participate voluntarily, including 14 participants as the tinnitus group, and 14 subjects as the control group. Tinnitus group inclusion criteria were (i) 18-60 years old, (ii) either unilateral or bilateral non-pulsatile tinnitus (> 3 months of duration), and (iii) audiogram thresholds ≤ 25 dB HL. All procedures were approved by the Institutional Scientific Ethic Committee of the Clinical Hospital of the Universidad de Chile (approval number OAIC 016/20042016), and all participants signed a written informed consent.

### Audiology

Hearing thresholds were measured using air-conduction pure-tone audiometry (AC40, Interacoustics, Denmark). We calculated the pure tone average (PTA) for each ear by averaging hearing thresholds at 0.5, 1, 2 and 4 kHz frequencies. To discard conductive hearing loss, we also measured the external ear canal acoustic immittance in both ears with a clinical device (AT235H, Interacoustics, Denmark).

### Behavioral task alternating between visual and auditory attention

Subjects performed the same behavioral paradigm described in Dragicevic et al., 2019, in which selective attention to visual and auditory modality alternated consecutively across trials. Briefly, as depicted in Figure 1, during visual attention trials, subjects had to focus their attention on the pointer of a clockwise-revolving clock (1 Hz), and report the offset time of a visual cue while ignoring DPOAE eliciting tones (f1 and f2). The onset of this visual cue (color change on the rim of the clock) indicated the period of focused visual attention (variable duration between 1.5 and 2.5 s) to the time indicated by the position of the clock’s pointer. After subject responses, the task switched to selective auditory attention, and the clock changed from a coherent, clockwise revolution to an incoherent, random positioning of the pointer without evoking a sense of motion. A brief gap of silence interrupted the DPOAE-eliciting tones appearing in a period of variable duration (between 1.5 and 2.5 s), and subjects had to report gap detection while looking at the center of the clock and ignoring the clock. After the behavioral response to the gap in DPOAE, the clock recovered its clockwise motion, and after a period of variable duration (between 2 and 2.5 s), the visual cue started. No silent gaps were presented during the visual task. Subjects gave their visual task response by first pressing a button, which triggered opposite and slower (1/3 Hz) rotation of the clock, and releasing the button at the desired position. Initial training of this response modality was given until motor error was minimized to about 3.6°, corresponding to the minimum angular step programmed in our display (360°/100).

**Figure 1.**
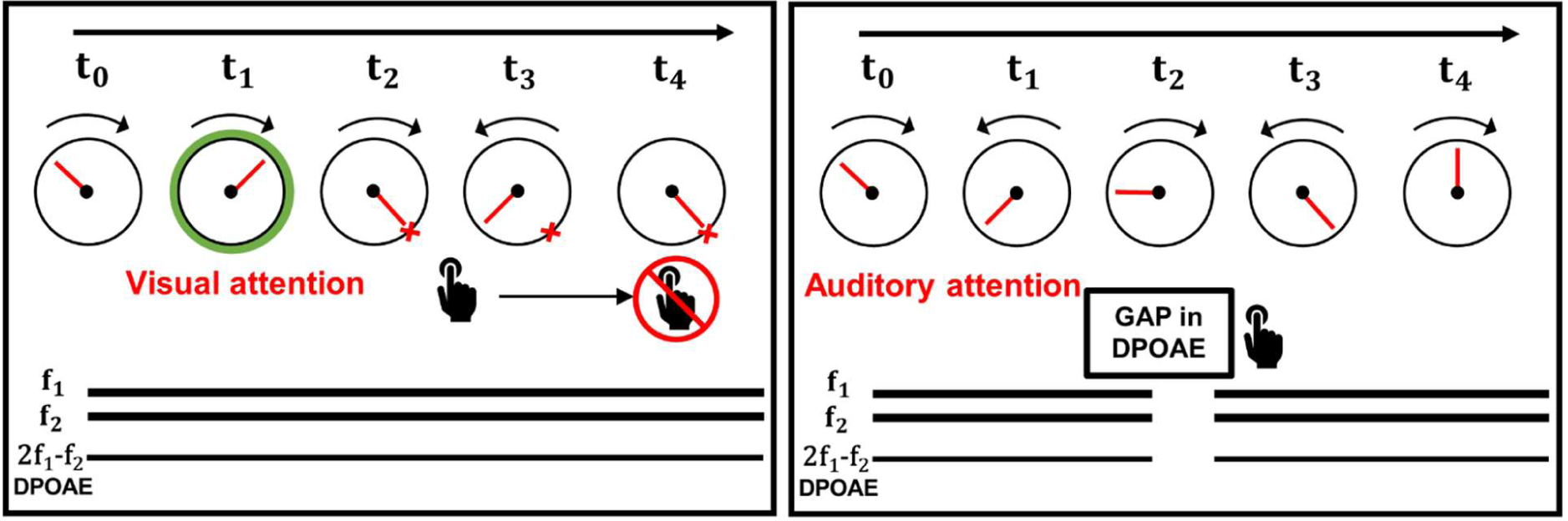
Behavioral task used to evaluate visual and auditory attention. A. Visual attention: Subjects had to report the time of the clock’s tick at the off-set of the green circle (visual attention cue). Simultaneously two tones (f1 and f2) that elicit DPOAEs were presented as auditory distractors. B. Auditory attention: Individuals had to respond to the gap of silence embedded in the f1 and f2 stimuli while the clock’s tick is moving randomly. DPOAE were recorded continuously during the whole task.

### Data acquisition system and general procedures

Continuous 32-channels EEG and 4-channels electro-oculogram (EOG) signals were recorded using Tucker-Davis Technologies hardware (PZ3 for EEG and RA4PA for EOG). Simultaneously, the DPOAE signal was recorded using an Etymotic Research microphone (ER10-C). EEG, EOG, and DPOAE signals were synchronized with a desktop computer running a custom software written in LabWindows/CVI 9.0 via a National Instruments A/D converter that controlled visual stimuli on a high refresh rate screen (100 Hz) and auditory stimuli by TTL pulses. Volunteers were prepared for EEG and DPOAE recordings, including scalp cleaning with alcohol, inspection of the external ear canal to assess the presence of earwax, selection and calibration of f1 and f2 primary tones to elicit DPOAE. In addition, training blocks were presented before data acquisition to ensure the full comprehension of the behavioral task (see Dragicevic et al., 2019 for more details).

### Data pre-processing

EEG and DPOAE single trial channels were evaluated by means of visual inspection. We carried out the rejection of trials considering the amplitude of the complete DPOAE signal for each recording block, this was done through cursors that delimited the allowed amplitudes. The trials that passed the described visual inspection process were individually inspected using the ELAN software (Aguera et al., 2011). For this procedure, the DPOAE channels, vertical EOG, horizontal EOG, and the EEG channels Fz, F3, Fc1, Cz, Fc2, F4, P3, O1, Pz, O2 and P4 were considered. Additionally, we applied the independent component analysis (ICA) technique to EEG channels to remove artifacts related to blinking and cardiac rhythm using the EEGLAB toolbox (v. 14.1.2) for Matlab.

### DPOAE virtual channel

We generated a DPOAE ‘virtual’ channel using a Hilbert-based approach to obtain the oscillation amplitude of the 2f1-f2 DPOAE component in the 1-50 Hz band. The method uses a band-pass filter of the signal with an attenuation value > 100 dB and centered at the frequency of the DPOAE (2f1-f2). After that, the envelope of the filtered signal was calculated using the Hilbert method. The signal envelope of the 2f1-f2 DPOAE amplitude was subsequently processed in the frequency domain in conjunction with EEG channels.

### Data analysis

For the EEG and DPOAE channels, time and frequency domain averages across trials were calculated, we used time windows of ± 1,500 ms aligned to the onset of visual or auditory attention periods. In the case of the visual modality, the trials were locked to the onset of the visual cue (0 ms for visual attention) that marked the beginning of the visual selective attention period. For the auditory modality, the onset of the selective attention period (0 ms for auditory attention) was determined by the end of the visual period.

Spectrograms were analyzed between 1 and 50 Hz at 1 Hz resolution. For each subject and channel, the spectrum of the single trials was obtained using Morlet wavelets. Then, frequency-specific z-scores were obtained based on each trial baseline (−1500 to 0 ms); that is, for each trial and frequency, the mean and standard deviation of the baseline was calculated, and the resulting spectrogram between 1 and 50 Hz was represented in z-scores. Finally, the spectrograms were averaged for each subject.

To evaluate possible significant differences in time spectrograms between auditory and visual attention in control and tinnitus groups, we performed permutation tests (Maris & Oostenveld, 2007) using a within-group approach. The general procedure consisted of computing the subtraction between the time-frequency matrices represented as z-scores. Then, a mask generated by the permutations test (from the two groups of n=14) was performed to identify significant differences in the differential spectrograms. We used a threshold of 0.05 for significance, 16,000 permutations, a maximum number of clusters of 3, and the two-sided option for detecting negative and positive clusters.

The differences in the time courses of the cochlear and electroencephalographic oscillations were evaluated by implementing a slope-to-peak analysis. This was done for each subject by calculating the point-to-point derivatives of the means of the z-score spectrograms in the 1-8 Hz band during the auditory and visual attention periods. The derivatives were calculated from 0 ms to the point at which it changed sign, then these values were averaged. Then, individual values were averaged for each channel (by modality and group). The statistically significant differences in the mean slope values were assessed with the Mann-Whitney test. In all the procedures mentioned above, the EEG channels chosen as relevant for the detailed analysis were Cz, Fz, and O1. These channels were selected as a measure of oscillatory activity in the auditory cortex, prefrontal cortex, and visual cortex, respectively that were also used in Dragicevic et al.,2019.

### Angular response analysis of the visual selective attention task

The behavioral responses in the visual period were recorded as the angular deviation relative to the correct answer, independent of the absolute target angle, such that a perfect response corresponded to 0° of deviation. We analyzed the angular responses using the CircStat Toolbox for Circular Statistics (Berens, P. 2009). Statistical comparisons between tinnitus and control groups were performed using the Watson-Willis test.

## Results

We recorded simultaneous EEG and DPOAE signals in 28 subjects, including 14 individuals with tinnitus (mean age 38.1 ± 9.1 years (mean ± SD)), and 14 controls (mean age 34.2 ± 8.0 years; p=0.12, t-test) while they were performing a behavioral task that alternates between visual and auditory attention (Dragicevic et al. 2019). There were non-significant differences in hearing thresholds between tinnitus and control groups as evaluated by audiograms obtained from 0.125 to 16 kHz (Figure 2). The PTA (0.5 to 4 kHz) for the tinnitus group was: 9.20 ± 0.28 dB HL (mean ± SD), while for the control group was: 9.02 ± 0.30 dB HL, p=0,45, t-test). Regarding anxiety levels, there were non-significant differences between tinnitus and control groups, as evaluated by the State-Trait Anxiety Inventory (STAI) scores (STAI tinnitus: 37.1 ± 9.3; STAI controls: 34.4 ± 12.0; p=0.25, t-test). The mean Tinnitus Handicap Inventory (THI) for the group of tinnitus patients was 31.1 ± 19.7 points, ranging between 8 and 72 points. Table 1 summarizes individual demographic and audiological data from the 14 tinnitus and 14 controls included in this study.

**Figure 2.**
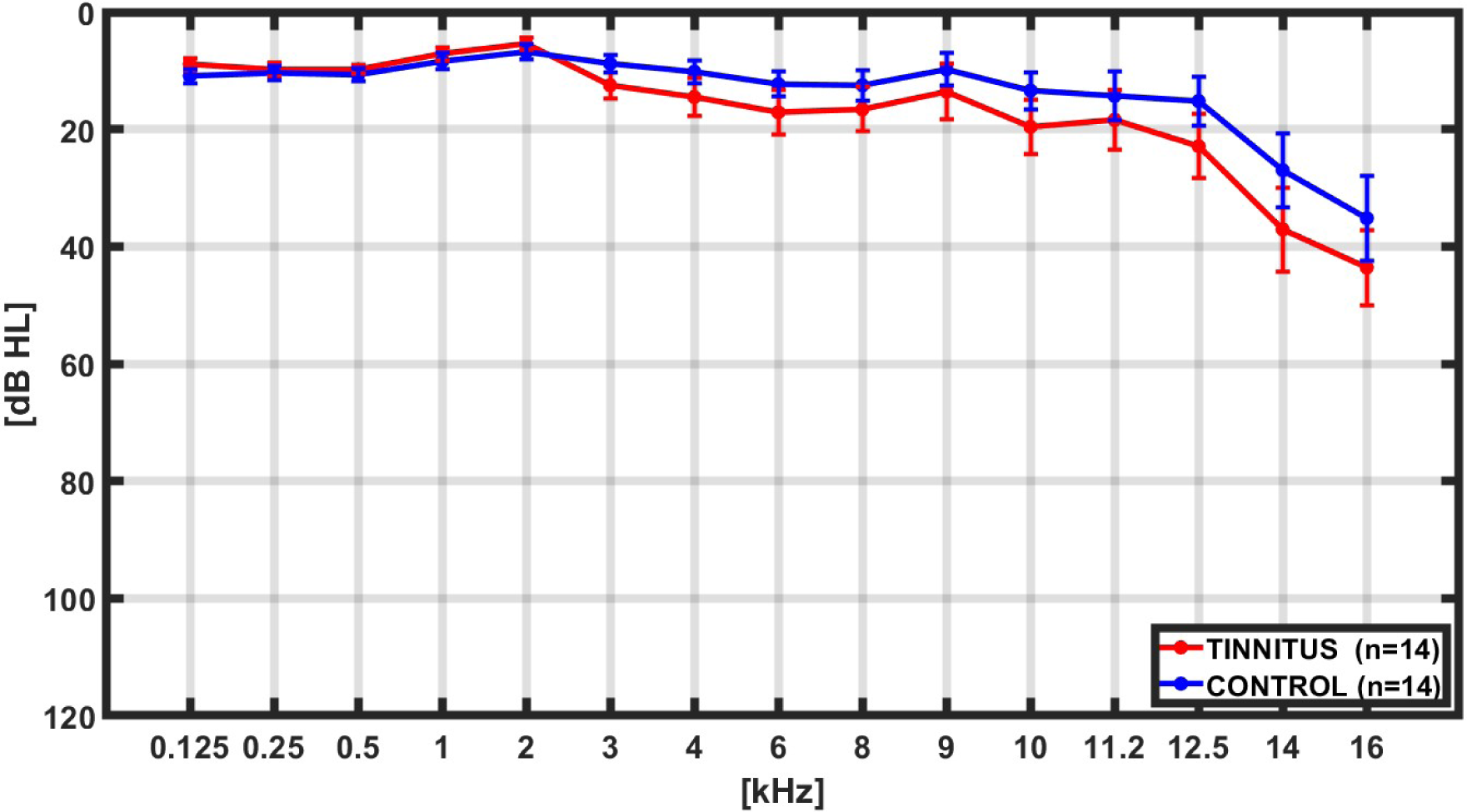
Grand average hearing thresholds. Bilateral average audiogram thresholds from 0.125 kHz to 16 kHz in controls (blue, n=14) and tinnitus (red, n=14) subjects. There were non-significant differences between groups in any of the evaluated frequencies.

**Table 1.** Summary of individual demographic, audiological and behavioral data in control (n=14) and tinnitus (n=14) subjects.

| Subject |  | Age | Sex | PTA<br>[dB HL] | Lat. Aud<br>[s] | LT. Vis<br>[s] | STAI | THI |
| --- | --- | --- | --- | --- | --- | --- | --- | --- |
| Tinnitus | 1 | 41 | M | 8.75 | 0.205 | 0.392 | 30 | 30 |
|  | 2 | 31 | M | 3.75 | 0.235 | 0.402 | 31 | 26 |
|  | 3 | 24 | F | 3.75 | 0.392 | 0.417 | 42 | 26 |
|  | 4 | 46 | M | 14.38 | 0.260 | 0.318 | 29 | 38 |
|  | 5 | 27 | M | 12.50 | 0.362 | 0.391 | 34 | 12 |
|  | 6 | 42 | M | 5.63 | 0.282 | 0.422 | 34 | 24 |
|  | 7 | 45 | F | 6.25 | 0.240 | 0.330 | 31 | 8 |
|  | 8 | 51 | M | 15.63 | 0.344 | 0.408 | 33 | 10 |
|  | 9 | 36 | M | 10.00 | 0.312 | 0.475 | 46 | 52 |
|  | 10 | 42 | M | 15.00 | 0.407 | 0.346 | 35 | 42 |
|  | 11 | 51 | F | 10.00 | 0.330 | 0.695 | 49 | 60 |
|  | 12 | 24 | M | 9.38 | 0.221 | 0.368 | 36 | 26 |
|  | 13 | 39 | M | 6.25 | 0.226 | 0.402 | 61 | 72 |
|  | 14 | 35 | M | 7.50 | 0.268 | 0.398 | 29 | 10 |
| Mean | - | 38.14 | - | 9.20 | 0.292 | 0.412 | 37.1 | 31.1 |
| Control | 1 | 42 | F | 9.38 | 0.387 | 0.540 | 24 | - |
|  | 2 | 24 | M | 3.75 | 0.281 | 0.345 | 29 | - |
|  | 3 | 28 | M | 11.88 | 0.224 | 0.376 | 29 | - |
|  | 4 | 48 | M | 18.13 | 0.351 | 0.357 | 22 | - |
|  | 5 | 31 | F | 6.25 | 0.215 | 0.390 | 24 | - |
|  | 6 | 42 | F | 8.13 | 0.389 | 0.497 | 32 | - |
|  | 7 | 25 | F | 7.50 | 0.404 | 0.510 | 68 | - |
|  | 8 | 41 | M | 7.50 | 0.248 | 0.339 | 32 | - |
|  | 9 | 31 | M | 16.25 | 0.329 | 0.420 | 28 | - |
|  | 10 | 45 | M | 8.75 | 0.409 | 0.503 | 43 | - |
|  | 11 | 34 | F | 11.88 | 0.239 | 0.348 | 28 | - |
|  | 12 | 32 | M | 5.63 | 0.274 | 0.351 | 38 | - |
|  | 13 | 25 | M | 6.88 | 0.276 | 0.345 | 46 | - |
|  | 14 | 31 | M | 4.38 | 0.451 | 0.485 | 38 | - |
| Mean | - | 34.21 | - | 9.02 | 0.320 | 0.415 | 34.4 | - |

### Behavioral responses

In relation to the behavioral performance, the mean response latency to the auditory target (silent gap in DPOAE) was shorter (306 ± 73 ms) than the mean response latency to the visual cue (413 ± 82 ms; p<0.01, t-test). When comparing the behavioral performance between tinnitus and control groups, we found non-significant differences in the mean response latencies to the visual and auditory targets (tinnitus (auditory task: 292 ± 46 ms; visual task: 412 ± 33 ms) and control groups (320 ± 40 ms (p=0.32, t-test); visual task: 415 ± 33 ms (p=0.92, t-test))(Table 1, Figure 3).

**Figure 3.**
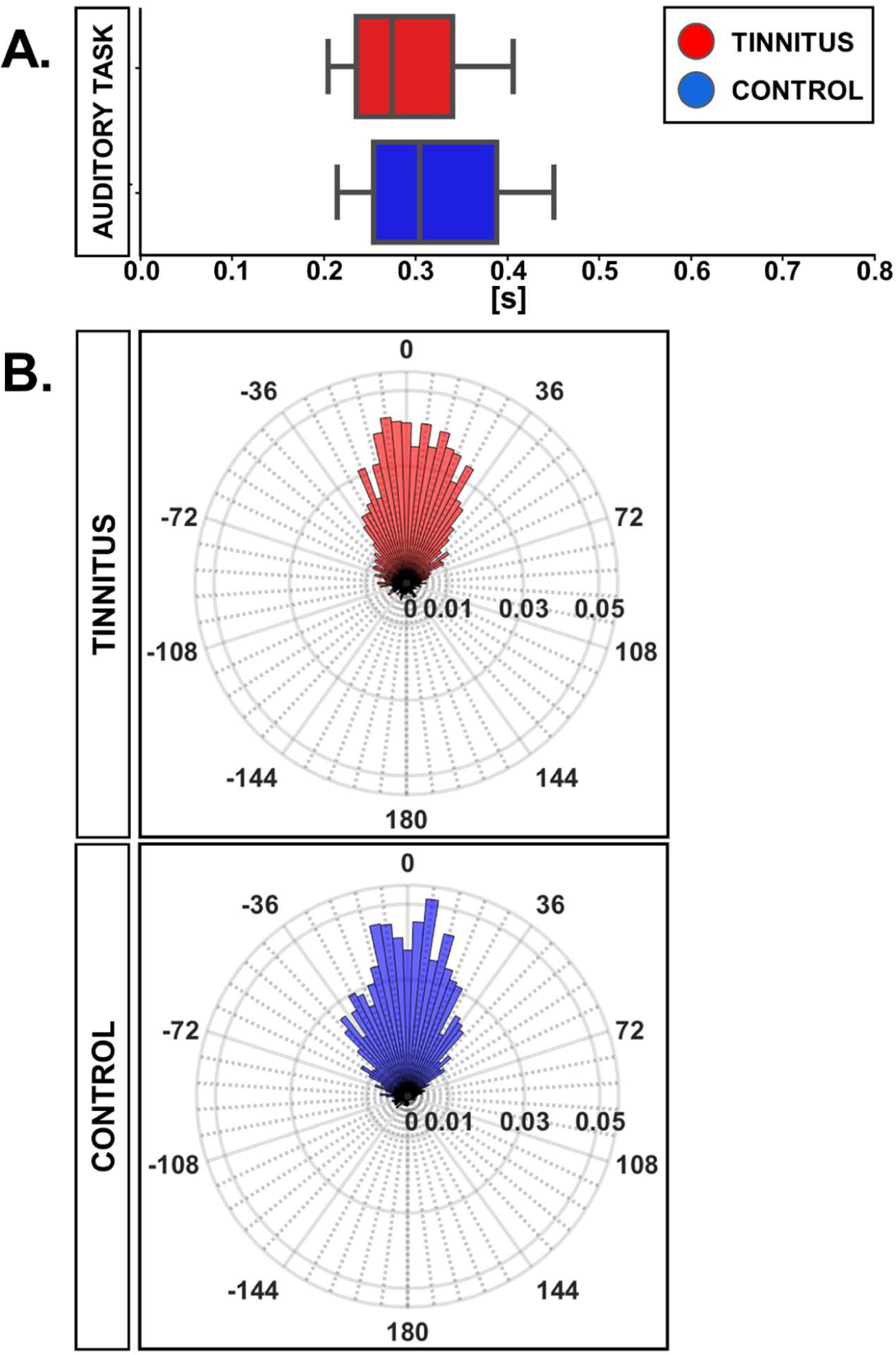
Histogram distribution of the latencies of correct behavioral responses to targets during the visual and auditory attention task in control and tinnitus groups. (A) Box-plots showing the latencies of responses to the gap in DPOAE during the auditory task in control and tinnitus groups. (B) Circular histograms of behavioral responses during the visual attention task, in which 0° represents the clock’s tick target position. The magnitude of the column bars illustrates the probability of a given response position in the clock referenced to the visual target offset.

We performed circular analyses statistics of the behavioral responses during the visual selective attention period (angular responses). The 14 subjects of the tinnitus group contributed with 2388 responses, while 14 individuals of the control group provided 2383 angular responses. The mean angle for the tinnitus group was 0.98° with a confidence interval (CI-95%) of [−1.10°, 3.07°], while for the control group was −5.03° with CI-95% of [−6.79°,−3.26°]. The Watson-Willis test, a circular analog of the two-sample t-test shows that the differences in angular responses between controls and tinnitus groups was significant (p=6.24 x 10^−6^).

The resultant vector length of the circle distribution of behavioral responses in the visual task can be used as a measure of dispersion (of value 1 when all responses are associated with the same angle), being 0.68 for the tinnitus group and 0.77 for the controls. We used the Von Mises distribution (μ: location, the data is clustered around this value; κ: concentration, the reciprocal measure of dispersion) to fit data. The values for the tinnitus group were (μ_T_, κ_T_)=(0.98°, 1.90), and for controls (μ_C_, κ_C_)=(−5.03°, 2.57). κ_T_ < κ_C_ indicates that the control group data are more concentrated around μ_C,_ and therefore we can conclude that the individuals in the tinnitus group presented less precise responses than controls in the visual task. The normalized polar histograms for both groups can be found in Figure 3B.

### Oscillatory activity

Next, we evaluated the oscillatory activity in EEG and DPOAE signals during auditory and visual attention periods in the control and tinnitus groups. Similar to our previous work (Dragicevic et al., 2019), we selected four regions of interest (ROIs), including Cz, Fz, O1, and DPOAE channels for frequency analyses. In agreement with previous works (Dragicevic et al., 2019; Kohler et al., 2021; Gemmacher et al., 2022), the grand average of the EEG and DPOAE spectra confirmed the presence of low-frequency oscillations in theta and delta frequency bands (<10 Hz) during the auditory and visual attention periods in the control and tinnitus groups (Figure 4).

**Figure 4.**
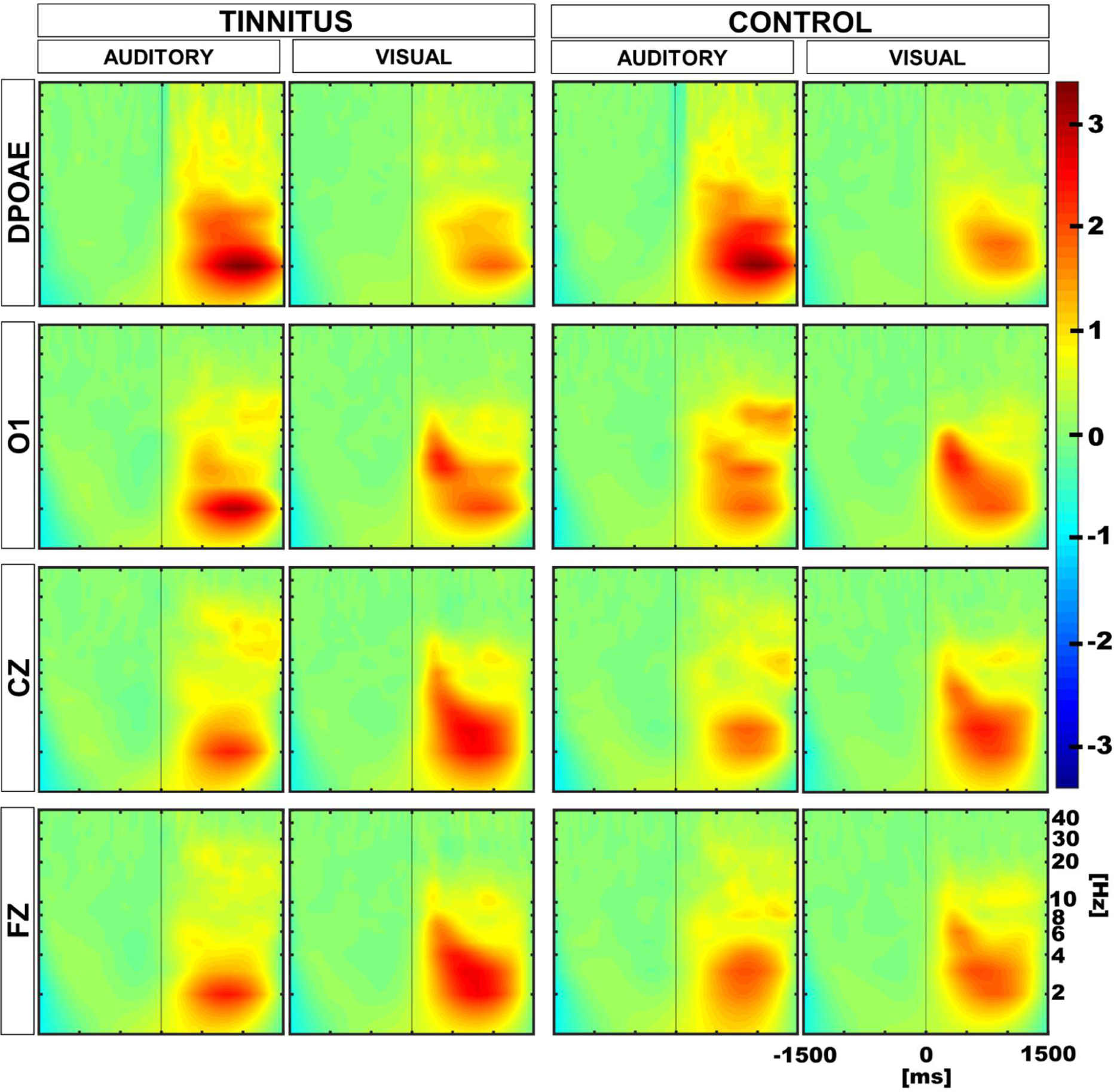
Grand average time spectra calculated for control and tinnitus groups during visual and auditory attention tasks. The color intensity scale was normalized to z-scores in all conditions. The vertical line in the panels illustrate time “0” in auditory and visual tasks. Note the presence of low-frequency oscillatory activity in theta and delta bands (<10 Hz) in EEG and DPOAE channels during the selective attention period (from 0 to 1500 ms). A significant reduction in the magnitude of the oscillatory activity in the cochlear channel (DPOAE) was observed in control and tinnitus groups during the visual attention period as compared to the auditory attention period. An increase in the theta power is observed in the Fz channel during the visual attention period in tinnitus.

### Oscillatory power in the 1-8 Hz band in the auditory versus visual attention periods

To evaluate differences in the oscillatory power of the delta and theta frequency bands in the ROIs between the auditory and visual attention periods, we considered the auditory and visual trials in the 1-8 Hz frequency band in the period between 200 and 1500 ms after the cue onset. Performing a paired t-test, we found a significant reduction of the oscillatory activity in the DPOAE channel during the visual attention period in the control and tinnitus groups, compared to the oscillatory activity in the auditory attention period (DPOAE-tinnitus: visual: 0.632, auditory: 1.185, p=0.0032; DPOAE-control: visual: 0.638, auditory: 1.255, p=0.0283; values on Z-score). Using the same criteria, we found a significant increase in the oscillatory power during the visual period in the Fz EEG channel for the tinnitus group (Fz-tinnitus: visual: 1.030, auditory: 0.688, p=0.0302; values on Z-score), but not for the control group (Fz-control: visual: 0.827, auditory: 0.815, p= 0.4775). In addition, we found significant increases in the oscillatory power in Cz channels for the visual attention period in both groups (Cz-tinnitus: visual: 1.051, auditory: 0.688, p=0.0171; Cz-control: visual: 0.940, auditory: 0.619, p=0.0087; values on Z-score). There were non-significant differences between visual and auditory oscillatory amplitudes in the occipital channel in both groups (O1-tinnitus: visual: 0.878, auditory: 0.834, p=0.3775; O1-control: visual: 0.847, auditory: 0.875, p=0.3849; values on Z-score).

### Oscillatory power in the 1-8 Hz band between control and tinnitus

Afterwards, we compared group differences between tinnitus and controls. We found a significant difference in the Fz channel for the visual task with a higher value for the tinnitus group (visual: Fz-tinnitus: 1.030, Fz-control: 0.827, p=0.040; values on Z-score). The other comparisons between ROIs in tinnitus and control groups were not significant.

### Permutation tests to compare individual differences

Next, to evaluate whether the differences observed in the amplitude of DPOAE and EEG low-frequency oscillations during the auditory and visual attention periods were different in tinnitus and controls at the individual level, we performed permutation tests using a within-subjects approach. Figure 5 shows the results of these permutation tests, uncorrected and corrected for multiple comparisons. Note that after multiple comparison correction, there is a significant positive cluster in the delta band in the DPOAE channel of the tinnitus individuals, illustrating that the suppression of the DPOAE oscillatory activity during visual attention in the delta band is stronger in tinnitus. We also found a significant negative cluster in the theta band in the Cz channel of the control group.

**Figure 5.**
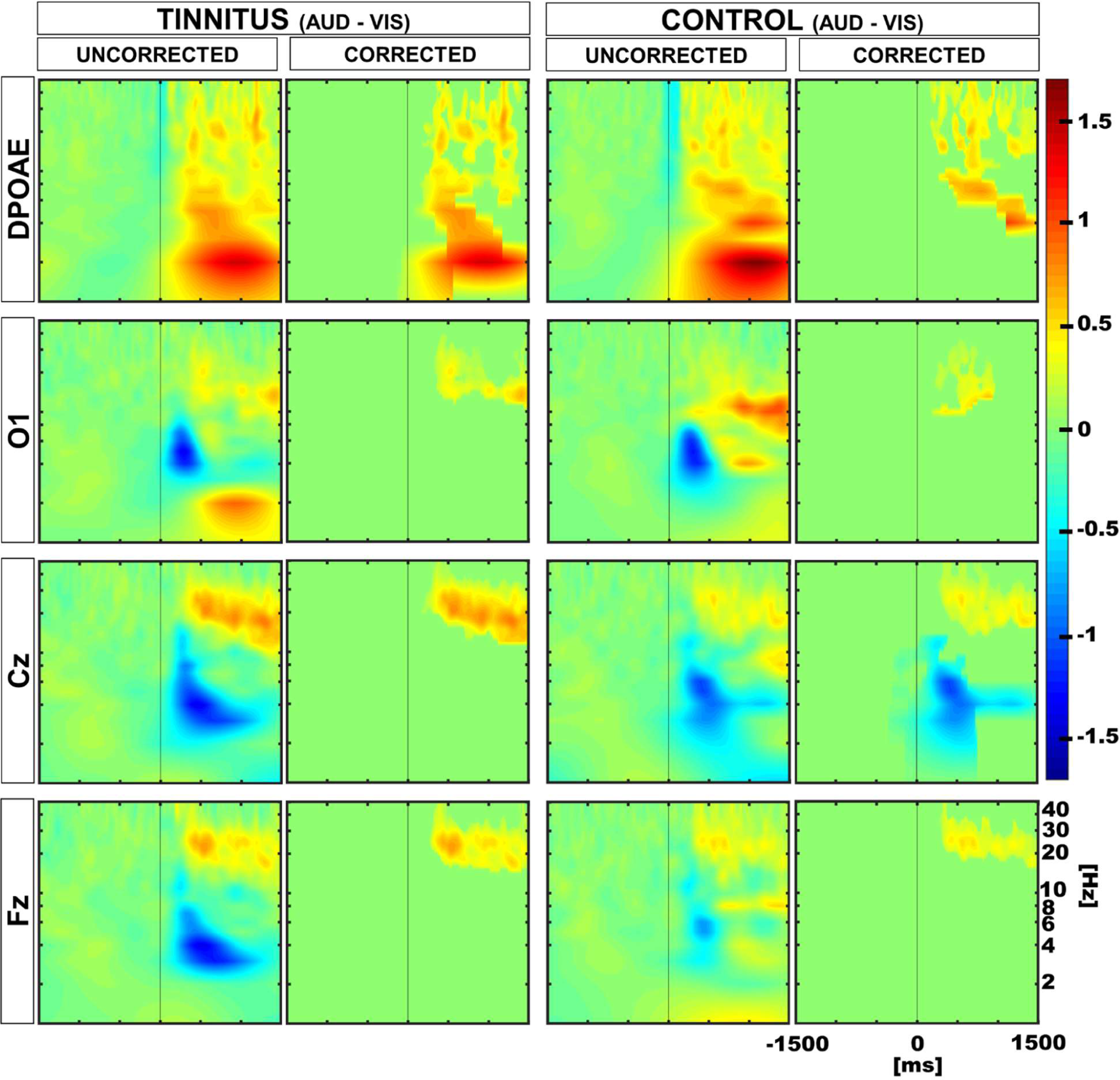
Time spectra average differences between auditory and visual periods using permutation tests in a within subjects approach. Color represents the grand average of the differences in Z-score. Red and blue clusters represent significant differences in the permutation test, while non-significant differences are shown in green. The first and third column show uncorrected results, while the second and fourth illustrate multiple comparison corrected results. Positive significant differences are represented in red (i.e. auditory > visual), while negative differences are illustrated in blue (i.e. auditory < visual). Note that after correction, there is a significant positive cluster in the delta-theta band in the DPOAE channel of the tinnitus group, and a significant negative cluster in the theta band in the Cz channel of the control group.

### Time course of the oscillatory changes in the 1-8 Hz frequency band

Finally, we explored the temporal course of the oscillatory power of the 1-8 Hz band in the auditory and visual attentional periods, in control and tinnitus groups. Figure 6 displays the grand averages of the temporal course of the power increase in the 1-8 Hz band (mean ± SEM, 14 tinnitus, 14 controls for each modality) for the DPOAE, O1, Cz, and Fz channels. To assess the differences between the temporal profiles of each channel, we performed a curve slope analysis, computing the mean slope values between 0 ms and the point where the slope began to decrease. The analysis showed that for the auditory modality, the mean slopes were steeper for the DPOAE channel in both groups compared to the mean slope for the EEG channels (tinnitus mean slope and p-value for DPOAE vs. EEG channel: DPOAE: 0.0023, O1: 0.0017 (p=0.162), Cz: 0.0014 (p=0.047), Fz: 0.0013 (p=0.015); control mean slope and p-value for DPOAE vs. EEG channel: DPOAE: 0.0026, O1: 0.0017 (p=0.0183), Cz: 0.0014 (p=0.0007), Fz: 0.0016 (p=0.0031)). Regarding the visual modality, the mean DPOAE slope values were lower than the mean slope of EEG channels in both groups (tinnitus mean slope and p-value for DPOAE vs. EEG channel: DPOAE: 0.0011, O1: 0.0035 (p=0.00031), Cz: 0.0033 (p=0.00004), Fz: 0.0030 (p=0.00006); control mean slope and p-value for DPOAE vs. EEG channel: DPOAE: 0.0014, O1: 0.0036 (p=0.0003), Cz: 0.0027 (p=0.0013), Fz: 0.0023 (p=0.0348)).

**Figure 6.**
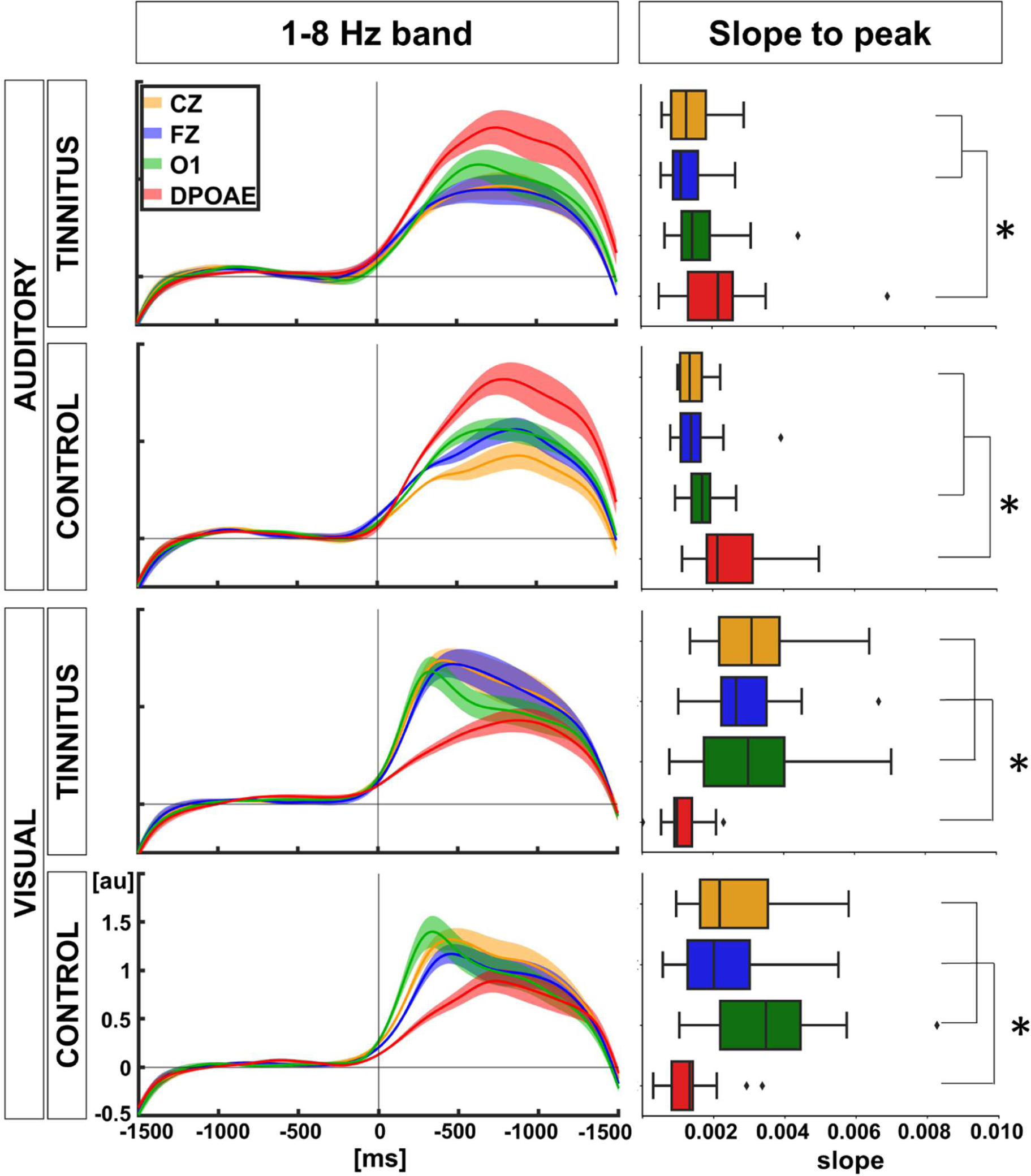
Temporal course of the amplitude of 1-8 Hz frequency band between EEG and DPOAE channels and comparison of slope values between individuals (*p<0.05)

Mean slope differences comparisons between tinnitus and control groups for visual and auditory modalities were non-significant (auditory: tinnitus vs. control: DPOAE: p=0.22, O1: p=0.27, Cz: p=0.47, Fz: p=0.13; visual: tinnitus vs. control: DPOAE: p=0.35, O1: p=0.30, Cz: p=0.16, Fz: p=0.10).

## Discussion

We found that cortical and cochlear oscillatory activity in the delta/theta bands were present in the tinnitus group, and similar to the control group, there was a significant decrease in DPOAE oscillatory amplitude during the visual attention period as compared to the auditory attention period, while frontal oscillatory activity (Fz) was increased during the visual attention task only in the tinnitus group. In addition, we found a significant cluster in the delta band in tinnitus when using within-group statistics to compare the difference between auditory and visual DPOAE oscillatory power. Finally, there was a significant difference in the temporal course of the oscillatory power ascending slope (in the 1-8 Hz band) when comparing the auditory and visual attention periods, but not between control and tinnitus groups.

### Behavioral differences between control and tinnitus

Behavioral comparisons between control and tinnitus yielded non-significant differences in the auditory task, and only subtle differences in visual attention. In the latter task, the mean angle values for both groups showed that subjects in the tinnitus group tended to report their responses before the clock pointer reached the target. In addition, the angular dispersion showed that the control group tended to respond more accurately than the tinnitus group. Together, these results might be explained by the known elevated anxiety levels of tinnitus sufferers (Kleinstäuber and Weise, 2020; Chen et al., 2023). The literature reports that tinnitus sufferers have psychological profiles showing more anxiety and with greater distractibility than the general population (Araneda et al., 2015; Kleinstäuber and Weise, 2020; Chen et al., 2023). However, unexpectedly, our analysis of the anxious traits (STAI) did not show significant differences between the two groups. This might be explained by the high level of anxiety observed in the general population, especially after the COVID-19 pandemia (Phalswal et al., 2023), which might have masked anxiety differences between control and tinnitus.

### Low-frequency cochlear oscillations are suppressed during visual attention

Previous works using DPOAE, external ear canal pressure and cochlear implant low frequency signals have evidenced the presence of low-frequency oscillations in the delta and theta bands at the most peripheral level of the auditory pathway during selective attention paradigms (Dragicevic et al., 2019; Kohler et al., 2021; Gemmacher et al., 2022; Köhler and Weisz 2023). Here, we used the same paradigm as Dragicevic et al., 2019, confirming the presence of these infrasonic oscillations in a new cohort of volunteers with and without tinnitus. Moreover, our data also confirmed a reliable suppressive effect of cochlear oscillatory activity during visual attention as compared to the auditory modality (Dragicevic et al., 2019; Kohler et al., 2021; Gemmacher et al., 2022). Altogether, these results show that the magnitude of cochlear oscillatory activity (delta and theta frequency bands) depends on the attentional modality, with increased amplitude during auditory attention and diminished amplitude with visual attention. Whether this effect is limited to the attentional mechanisms or is a more general cognitive mechanism is still an open question (Marcerano et al., 2021).

### Oscillatory power in the 1-8 Hz band in the between control and tinnitus

The next question was whether cochlear and cortical oscillatory activity were altered in tinnitus patients as compared to control patients during visual and auditory attention. We used two approaches, (i) a comparison between groups in the different conditions, and (ii) a second approach using within-subjects permutation tests for studying individual differences in tinnitus and control subjects between the auditory and visual attention periods.

All significant differences at the DPOAE level were found when using the within-subjects approach, while when using the group-average approach there were non-significant differences between controls and tinnitus. These findings indicate that the cochlear oscillatory activity is present in tinnitus and control groups, and that it might be a relatively stronger corticofugal activity in the tinnitus group that is affected by the inter-individual variability. Only when using an individual comparison (within subject approach), the difference between the cochlear oscillatory activity in the delta band was stronger for the tinnitus group (Figure 5, corrected DPOAE spectrograms). A speculative explanation for this higher corticofugal suppression of cochlear low-frequency oscillations in tinnitus could be the presence of a compensatory top-down mechanism for suppressing the phantom sound. Importantly, this finding was concomitant to an increase in the theta band in frontal regions, which has been found in more severe tinnitus (Czornik et al., 2022). Here we propose that the higher oscillatory activity in the theta band in the frontal region might indicate a greater compensatory top-down suppression to the cochlear receptor in the delta band during visual attention in tinnitus patients.

### Time course of the oscillatory changes in the 1-8 Hz frequency band

Finally, we studied the temporal course of the oscillatory amplitudes changes by measuring the differences in slopes along time of EEG and DPOAE oscillations. Our findings confirmed a temporal shift of the ascending phase of the cochlear oscillatory activity in the low frequency band with auditory and visual attention (Dragicevic, 2019), showing that during auditory attention, brain oscillations were preceded by the cochlear oscillations, while during visual attention, brain oscillatory activity precedes cochlear oscillations (Figure 6). These results are indicative of a peripheral correlate of the attentional switch at the delta-theta band that occurs in the transition between the auditory and visual modality. Importantly a similar neural correlate of attentional switching has been obtained by other groups in EEG recordings (Philips et al., 2014; Wang et al., 2016). On the other hand, when comparing the slopes of the temporal course of EEG and DPOAE low-frequency oscillations in tinnitus and control groups, we did not find significant differences, suggesting that the temporal dynamics of these oscillatory activity was not disrupted in tinnitus sufferers.

## Conclusions

We confirm the presence of infrasonic low-frequency cochlear oscillations in the delta and theta bands in tinnitus patients. In addition, we found that the corticofugal suppression of cochlear oscillations during visual attention is also present in tinnitus individuals, and as suggested by the within group analysis, it might be stronger in the delta band in tinnitus than in control subjects, and accompanied by an increase in the theta oscillatory power in frontal regions in tinnitus sufferers.

## Acknowledgments

This research was funded by ANID Proyecto Fondecyt Regular 1220607 to PHD, Fondecyt Postdoctorado 3230557 to VM, Fondecyt Postdoctorado 3200735 to CDD. Proyecto Basal ANID FB0008, Proyecto Milenio ICN09_015, and Fundación Guillermo Puelma to PHD. We would like to thank Fernando Vergara for his technical contributions.

